# SARS-CoV-2 Encodes a PPxY Late Domain Motif Known to Enhance Budding and Spread in Enveloped RNA Viruses

**DOI:** 10.1101/2020.04.20.052217

**Authors:** Halim Maaroufi

## Abstract

The current COVID-19 (<u>Co</u>rona<u>vi</u>rus <u>D</u>isease-20<u>19</u>) pandemic is affecting the health and/or socioeconomic welfare of almost everyone in the world. Finding vaccines and therapeutics is therefore urgent, but elucidation of the molecular mechanisms that allow some viruses to cross host species boundaries, becoming a threat to human health, must also be given close attention. Here, analysis of all proteins of SARS-CoV-2 revealed a unique PPxY Late (L) domain motif, ^25^PPAY^28^, in a spike (S) protein inside a predicted hot disordered loop subject to phosphorylation and binding. PPxY motifs in enveloped RNA viruses are known to recruit Nedd4 E3 ubiquitin ligases and ultimately the ESCRT complex to enhance virus budding and release, resulting in higher viral loads, hence facilitating new infections. Interestingly, proteins of SARS-CoV-1 do not feature PPxY motifs, which could explain why SARS-CoV-2 is more contagious than SARS-CoV-1. Should an experimental assessment of this hypothesis show that the PPxY motif plays the same role in SARS-CoV-2 as it does in other enveloped RNA viruses, this motif will become a promising target for the development of novel host-oriented antiviral therapeutics for preventing S proteins from recruiting Nedd4 E3 ubiquitin ligase partners.

## INTRODUCTION

The current COVID-19 (<u>Co</u>rona<u>vi</u>rus <u>D</u>isease-20<u>19</u>) pandemic caused by SARS-CoV-2 from Wuhan is affecting the health (both physical and psychological) and/or socioeconomic welfare of almost everyone in the world. Finding vaccines and therapeutics to fight SARS-CoV-2 is therefore urgent. In addition, it is imperative, now and in the future, to maintain research efforts aimed at elucidating the molecular mechanisms (tropism, cell entry, multiplication and spread, etc.) used by emerging viruses such as SARS-CoV-2 to infect humans. An improved understanding of these mechanisms should be viewed as an investment in our future to prevent and/or control the emergence of dangerous viruses that could have as or more severe consequences for humanity than SARS-CoV-2. The family *Coronaviridae* (CoVs) is large and comprises enveloped, positive-sense, single-stranded RNA viruses (Graham and Baric, 2010; Li, 2013). Among all RNA viruses, CoVs possess the largest genomes, ranging in size between 27 and 32 kb. The subfamily *Coronavirinae* comprises four genera, *Alpha-, Beta-, Delta-* and *Gammacoronavirus*, which can infect both mammalian and avian species. Phylogenetically, SARS-CoV-2 is more closely related to SARS-CoV-1 than to MERS-CoV, but all three are betacoronaviruses (betaCoVs) (de Wit et al., 2016).

The envelope-anchored SARS-CoV-2 spike (S) is a multifunctional glycoprotein with 1273 amino acid residues. Its structure was solved as a homotrimeric molecule with several domains. The S protein is subdivided into S1 and S2 subunits. S1 is formed by two domains, an N-terminal domain (NTD) and a C-terminal receptor-binding domain (RBD) that binds specifically to the host receptor angiotensin-converting enzyme 2 (ACE2) on the host cell surface (Li, 2012). The S2 subunit, which is also multidomain, induces viral-host membrane fusion.

The viral proline-rich Pro–Pro-x-Tyr (PPxY) Late or L-domain motif (Chen and Sudol, 1995) interacts with host proteins containing the WW-domain. The term “Late” reflects a late function in the virus life cycle (Garnier et al., 1996; Freed, 2002). L-domain motifs are identified in enveloped RNA viruses such as in the Gag protein of a number of retroviruses and in the matrix of arenaviruses, rhabdoviruses and filoviruses (Baillet et al., 2019; Bieniasz, 2006; Wirblich et al., 2008). The WW-domain is a small domain of 35–40 amino acids, named after two conserved tryptophan (W) residues in the sequence (Bork and Sudol, 1994). It is a fold of three-stranded antiparallel β-sheets that contains two ligand-binding grooves (Huang et al., 2000; Kanelis et al., 2001). RNA viruses use PPxY L-domain motifs to recruit specific WW-domain of host proteins of either Nedd4 (<u>N</u>euronal precursor cell-<u>E</u>xpressed <u>D</u>evelopmentally <u>D</u>own-regulated 4), or Nedd4 family members to facilitate their egress and spread (Bieniasz, 2006). Nedd4 family proteins have three domains, N-terminal lipid binding (C2) domain; WW modules, present in multiple copies; and a C-terminal Hect (for homologous to E6-associated protein C terminus) domain that contains the ubiquitin ligase activity.

Previous and current investigations are focusing on understanding the first step that allows SARS-CoV-1 and SARS-CoV-2 to enter host cells. This research examines primarily the interaction between the C-terminal receptor-binding domain (RBD) of S proteins and the host receptor angiotensin-converting enzyme 2 (ACE2). Here, the molecular mechanism of the last step (late function virus life cycle) of budding and release of virus is presented. The N-terminal of SARS-CoV-2 spike protein contains a PPxY L-domain motif that is known to hijack host WW-domain of Nedd4 E3 ubiquitin ligases and ultimately the ESCRT (Endosomal Sorting Complex Required for Transport) complex to enhance virus budding and spread. Importantly, this motif is absent in SARS-CoV-1, which could explain why SARS-CoV-2 is more contagious than SARS-CoV-1. Development of a novel host-oriented inhibitor targeting molecular interactions between PPxY and the host WW-domain holds potential in the fight against SARS-CoV-2 infection.

## RESULTS AND DISCUSSION

### The N-terminal of S protein contains a PPxY L-domain motif

Sequence analysis of SARS-CoV-2 spike protein using the Eukaryotic Linear Motif (ELM) resource (http://elm.eu.org/) revealed a short linear motif (SLiMs) known as PPxY Late (L-) domain motif, ^25^PPAY^28^ in the N-terminus of the S1 subunit. The PPxY L-domain motif is known to interact with proteins containing WW-domain(s). L-domain motifs have been identified in single-stranded enveloped RNA viruses (Table 1), including in the Gag proteins of several retroviruses and in the matrix of arenaviruses, rhabdoviruses and filoviruses (Baillet et al., 2019; Bieniasz, 2006; Wirblich et al., 2008). RNA viruses use PPxY L-domain motifs to recruit specific WW-domains of host Nedd4 family members and ultimately the ESCRT complex to facilitate their budding and egress (Bieniasz, 2006). Interestingly, the SARS-CoV-2 S protein and Nedd4 can be S-palmitoylated, suggesting that they can localize to similar membrane subdomains (Petit et al., 2007; Gordon et al., 2020; Chesarino et al., 2014) and may therefore be able to interact. The ScanProsite tool (https://prosite.expasy.org/scanprosite/) showed that the PPxY motif is present in SARS-CoV-2 but absent from SARS-CoV-1 proteins. It has been reported in enveloped RNA viruses that PPxY motif recruits Nedd4 E3 ubiquitin ligases and ultimately the ESCRT complex to enhance virus budding and release that means a high viral load, thus facilitating new infections. This suggests in SARS-CoV-2 that the PPxY motif, through its role in enhancing the viral load, could play a role in making SARS-CoV-2 more contagious than SARS-CoV-1 (To et al., 2020).

**Table 1.** PPxY late-domain motif in single stranded enveloped RNA viruses.

| Viruses | Family | disease in human | Geography | PPxY protein | Reference |
| --- | --- | --- | --- | --- | --- |
| HIV-1 | Retroviruses | AIDS | World | Gag | Strack et al., 2000 |
| HTLV-1 | Retroviruses | Aggressive lymphoma | World | Gag | Bouamr et al., 2003 |
| Rabies | Rhabdoviruses | Deadly brain inflammation | World | M | Harty et al., 2001 |
| Marburg | Filoviruses | Deadly hemorrhagic fever | West Africa | VP40 | Dolnik et al., 2010 |
| Ebola | Filoviruses | Deadly hemorrhagic fever | West Africa | VP40 | Harty et al., 2000 |
| Lassa virus | Arenaviruses | Deadly hemorrhagic fever | West Africa | Z matrix | Baillet et al., 2019 |

The PPxY motif is also present in the S protein of bat RaTG13 and pangolin GX-P5L, two SARS-related coronaviruses (Fig. 1A). Moreover, S proteins with PPxY motif are phylogenetically close (Fig. 1C). Interestingly, alignment of S protein sequences of two pangolin coronaviruses showed only two differences in the S1 subunit (amino acid at position 25 (P/Q) and deletion of ^448^GY^449^ in pangolin GX-P1E) (Lam et al., 2020; Zhang et al., 2020). In the future, it would be interesting to compare virulence of pangolin GX-P1E and pangolin GX-P5L coronaviruses to know if the loss of PPxY motif is important for virulence. Figure 1B shows differences among coronaviruses in nucleotide composition for the codon corresponding to Pro25 in SARS-CoV-2. Transition mutations (purine (A, G) to purine or pyrimidine (C, T) to pyrimidine mutations) are more likely than transversion (purine to pyrimidine or vice versa) mutations. Additionally, most substitutions occur in the third nucleotide of codons (silent mutations), less frequently in the first nucleotide, and the least frequently in the second nucleotide (Wang et al., 2007). Therefore, a mutation from Ala (<u>G</u>CT) to Pro (<u>C</u>CT) is more likely to occur than one from Gln (C<u>A</u>G) to Pro (C<u>C</u>G). Moreover, a double mutation Asn (<u>AA</u>T) to Proline (<u>CC</u>T) is even less probable (Fig. 1B).

**Figure 1.**
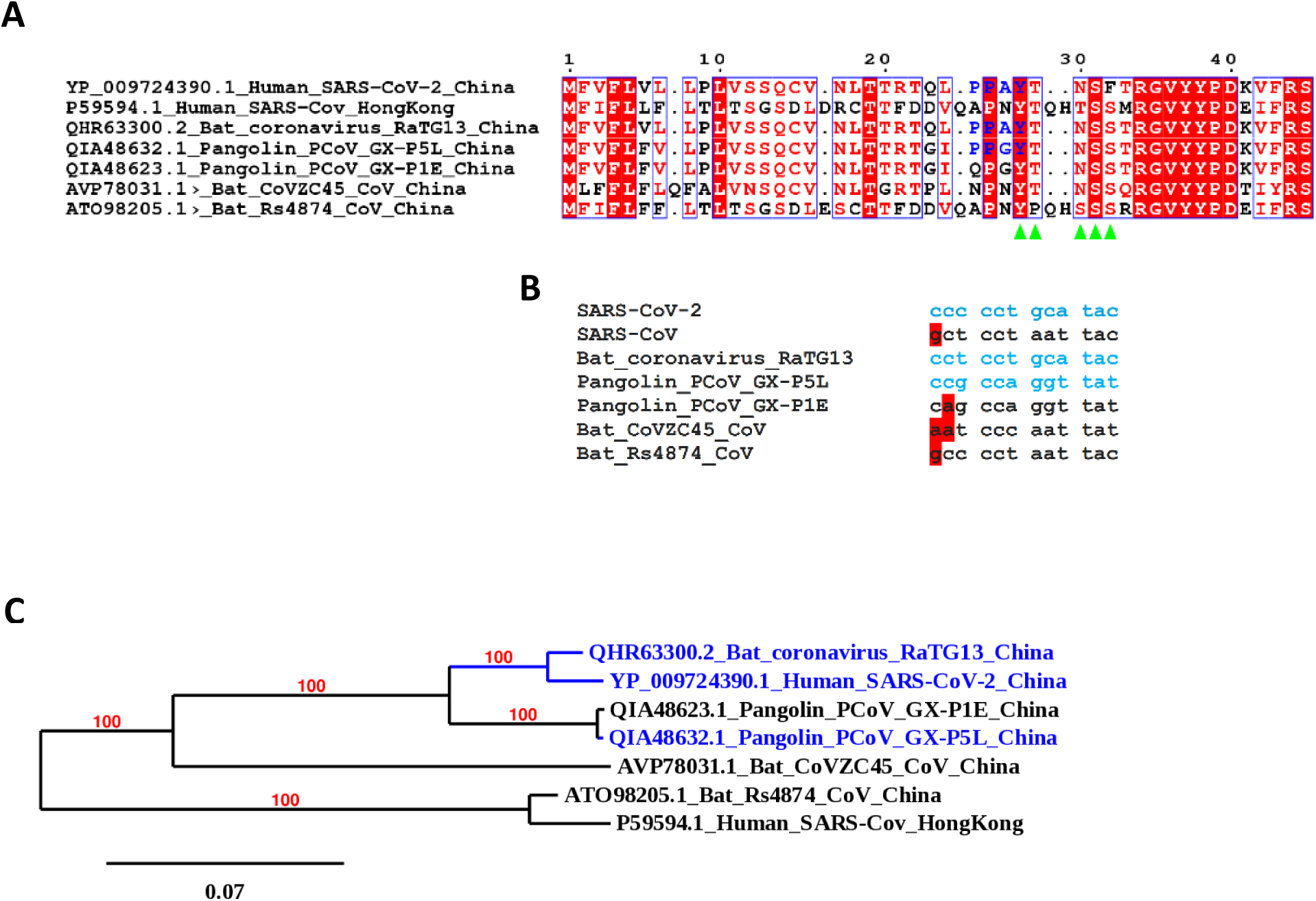
Multiple alignment of spike (S) protein of betacoronaviruses using the T-coffee software. **(A)** Localization of PPxY Late domain in N-terminal. PPxY Late domain motifs are colored in blue. The ^28^<u>Y</u>T<u>N</u>SF^32^ motif is indicated by green arrows. When its Tyr is phosphorylated, it is able to bind Src Homology 2 (SH2) domain. GenBank and UniProt accession numbers are indicated at the start of each sequence. The figure was prepared using ESPript (http://espript.ibcp.fr). (B) Alignment of codons of PPxY motifs. If nucleotides in red in codons could mutate into cytosine, they would produce proline residue. (C) Unrooted phylogenetic tree of S protein sequences of representative betacoronaviruses. The tree was constructed using Mr Bayes method based on the multiple alignment of complete S protein sequences by T-coffee. Betacoronaviruses in blue possess a PPxY motif. GenBank and UniProt accession numbers are showed at the start of each sequence.

### PPxY L-domain motif is in hot region

Scanning the SARS-CoV-2 S protein sequence with the DisEMBL software (Linding et al., 2003) revealed, in the N-terminal region, a unique hot disordered loop ^19^TTRTQL**PP**A**Y**TNSFT^33^ that includes the PPxY motif (Table 2), and this loop contains additional motifs. Indeed, the PPxY motif shares its tyrosine with an overlapping ^28^YTNSF^32^ motif (Fig. 1A) that enables binding to the SH2 domain of STAP1 when its Tyr28 is phosphorylated. STAP1 (Signal Adaptor Protein 1) is an adaptor protein with an SH2 and a PH class of lipid-binding domain (Wang et al., 1994). It is expressed in lymphoid cells and is phosphorylated by the Tec tyrosine kinase (Tec TK), which participates in B cell antigen receptor signaling (Ohya et al., 1999; Yang et al., 2001). The middle T-antigen of the polyoma virus, with its tyrosine residue in a phosphorylated state, has been shown to act as a binding site for the SH2 domain of the phosphatidylinositol-3’OH kinase 85K subunit (Dilworth et al. 1994). Another site, ^28^YTNS^31^ in the S protein, is also Tyr-phosphorylation-dependent for binding SH2 domain (Lin et al., 2006).

**Table 2.**
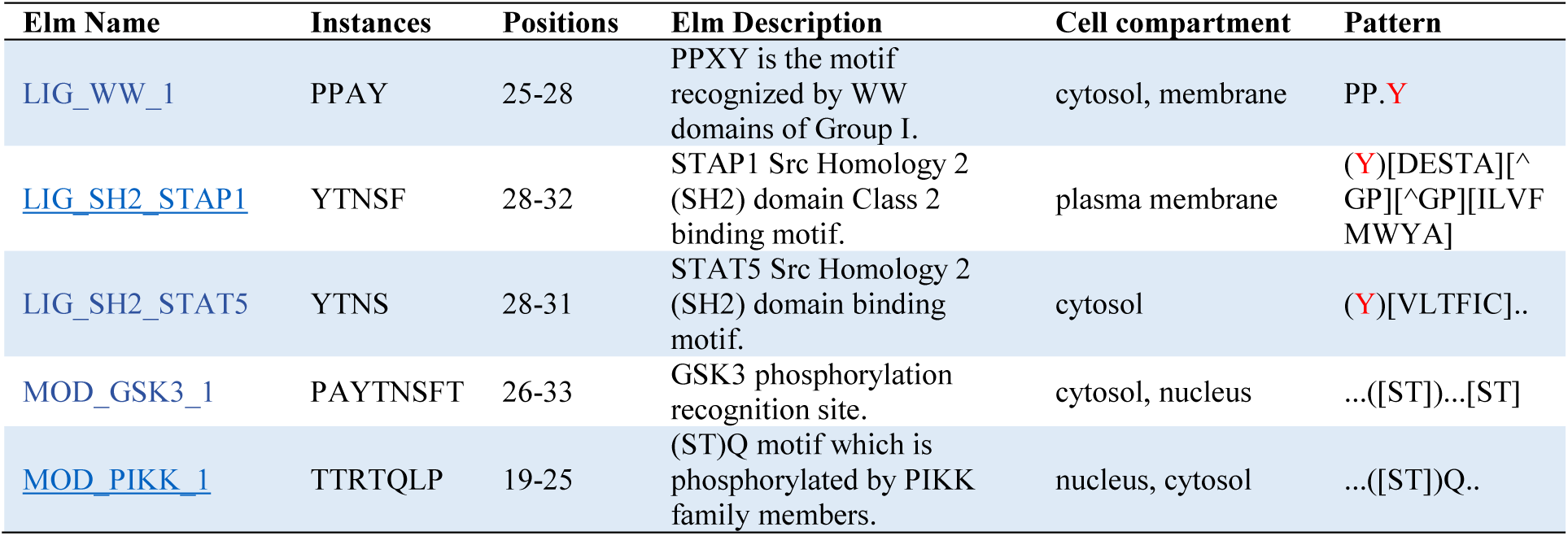
ELM motifs of hot disordered loop in SARS-CoV-2 S protein.

The hot region of the S protein also contains a Ser/Thr phosphorylated motif. Indeed, the PPxY motif shares its Pro25 with an overlapping ^19^TTR<u>TQ</u>L**P**^25^ motif that could be phosphorylated at TQ by a phosphatidylinositol 3- and/or 4-kinase (PIKKs). The Gln (Q) beside to the target Ser/Thr is critical for substrate recognition. PIKKs are atypical Ser/Thr kinases found only in eukaryotes (Imseng et al., 2108; Angira et al., 2020). Interestingly, Yang et al. (2012) showed that phosphatidylinositol 4-kinase IIIβ (PI4KB) is required for cellular entry by pseudoviruses bearing the SARS-CoV-1 S protein. This cell entry is highly inhibited by knockdown of PI4KB. Moreover, the same authors showed that PI4KB does not affect virus entry at the SARS-CoV-1 S-ACE2 interface. Finally, the motif ^26^PAY<u>**T**</u>NSF<u>**T**</u>^33^ is recognised by the Ser/Thr kinase GSK3 (Glycogen synthase kinase 3). Interestingly, the lipid kinase phosphatidylinositol 4-kinase II α (PI4KIIα) was reported to be a substrate of GSK3 (Robinson et al. 2014). This finding establishes a link with ^19^TTR<u>TQ</u>L**P**^25^motif, described above, that is predicted to be phosphorylated by GSK3.

In summary, proteins that interact with this hot disordered loop are connected to lipids (membrane), particularly with phosphatidylinositol, and bindings is regulated by phosphorylation. Recently, Besson et al. (2019) showed that the mitogen-activated protein kinase pathway and phosphatidylinositol metabolism are involved in the later stage of RABV infection.

### Interactions of PPxY L-domain motif with Nedd4-WW3-domain Ub ligase

To compare the interaction between the PPAY S protein and the Nedd4WW-domain with those of other PPxY-Nedd4WW-domain complexes, molecular docking was performed using the software AutoDock vina (Trott and Olson, 2010). Fig. 2 shows that the ^23^QL<u>**PP**</u>A<u>**Y**</u>TNS^31^ peptide of SARS-CoV-2 S protein is localized in the same region as ^112^TA<u>**PP**</u>E<u>**Y**</u>MEA^120^ motif of the Zaire Ebola virus Matrix protein VP40 (PDBid: 2KQ0), and important amino acid residues of the SARS-CoV-2 peptide are superposed with corresponding residues of the Ebola VP40 peptide, with the exception of different orientations for the aromatic ring of tyrosine (Fig. 2B and C). Iglesias-Bexiga et al. (2019) reported that the Nedd4-WW3 binding site is highly plastic and can accommodate PPxY-containing ligands with varying orientations. This structural superposition with PPxY of VP40 confirms the reliability of the AutoDock vina peptide docking module. Detailed knowledge of the molecular interaction between the ^23^QL<u>**PP**</u>A<u>**Y**</u>TNS^31^ motif and the Nedd4WW-domain will help design a peptide able to mimic the surface of the Nedd4 WW-domain and strongly bind the ^23^QL<u>**PP**</u>AYTNS^31^ peptide, thus preventing the recruitment of Nedd4 E3 ubiquitin ligase by the SARS-CoV-2 S protein (Zaidman and Wolfson, 2016). Indeed, Han et al. (2014) have developed two lead compounds that specifically block the VP40 PPxY-host Nedd4 interaction, displaying anti-budding activity in the nanomolar range.

**Figure 2.**
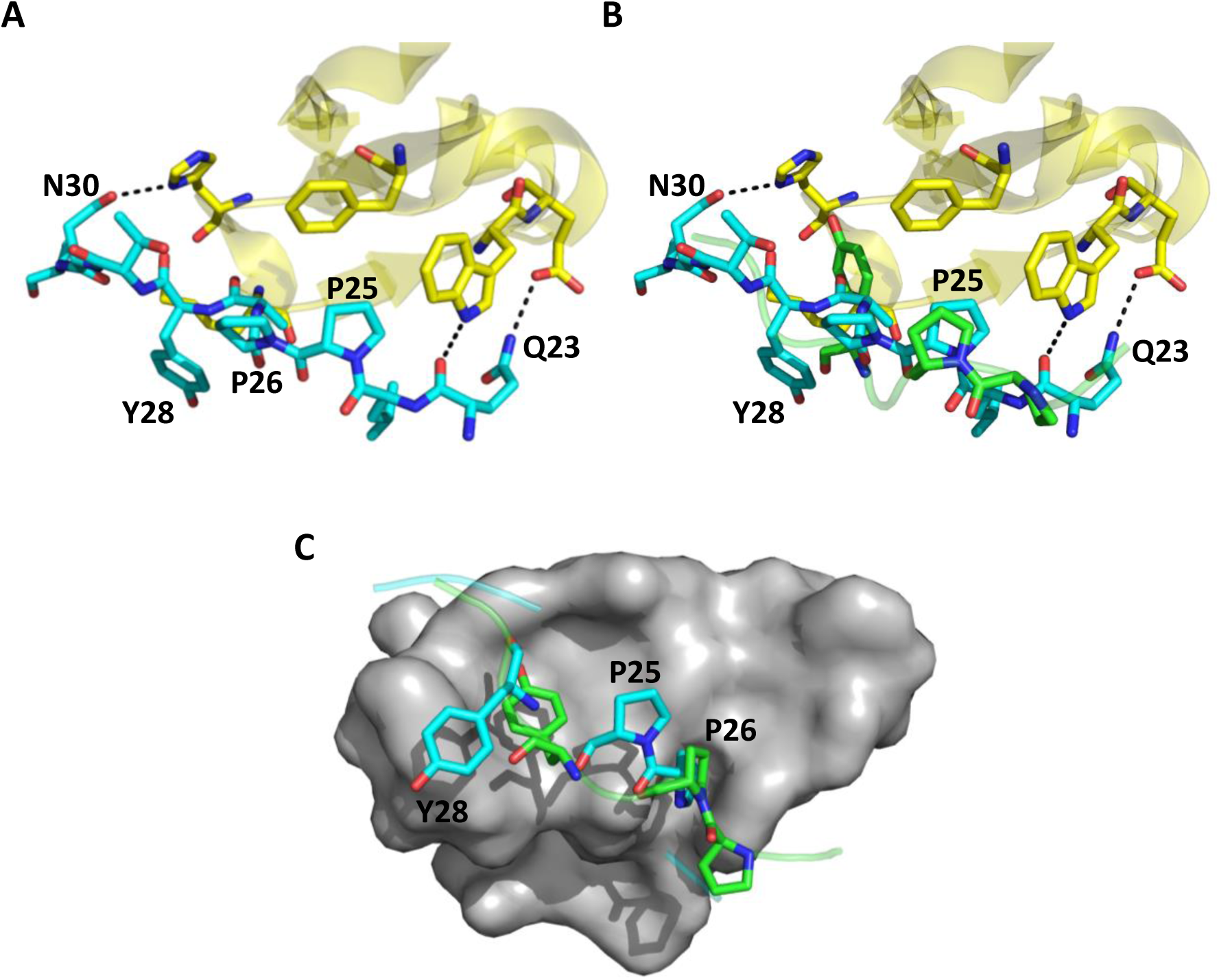
Cartoon and surface representation of human Nedd4 WW3-domain (PDBid: 2KQ0_A) with docked peptides. (A) ^23^QL<u>**PP**</u>A<u>**Y**</u>TNS^31^ peptide of SARS-CoV-2 S protein (cyan) and (B) ^23^QL<u>**PP**</u>A<u>**Y**</u>TNS^31^ superposed to Zaire Ebola virus Matrix protein VP40 (^112^TA<u>**PP**</u>E<u>**Y**</u>MEA^120^) peptide (green, PDBid: 2KQ0_B). (C) Surface representation of Nedd4 WW3-domain with peptides depicted as sticks. Images were generated using PyMol (www.pymol.org).

## MATERIALS AND METHODS

### Sequence analysis

To search probable short linear motifs (SLiMs), SARS-CoV-2 spike protein sequence was scanned using the eukaryotic linear motif (ELM) resource (http://elm.eu.org/). The PPxY L-domain motif was searched in proteins of SARS-CoV-2 and SARS-CoV-1 using https://prosite.expasy.org/scanprosite/

### 3D modeling and molecular docking

Unfortunately, the region containing the ^23^QL<u>**PP**</u>A<u>**Y**</u>TNS^31^peptide has not been resolved in any known structure of spike S protein. Therefore, the Pep-Fold (Thevenet et al., 2012) software was used to generate a *de novo* model of this peptide. Model quality of the peptide was assessed by analysis of a Ramachandran plot through PROCHECK (Vaguine et al., 1999).

Docking of the peptide into the human Nedd4 3rd WW-domain (PDBid: 2KQ0_A) was performed using the software AutoDock vina (Trott and Olson, 2010). The 3D complex containing the human Nedd4 3rd WW-domain and the peptide was refined using FlexPepDock (London et al., 2011), which allows full flexibility to the peptide and side-chain flexibility to the receptor.

### Phylogeny

Amino acid sequences of different spike (S) proteins, namely those of SARS-CoV-2 (accession: YP_009724390.1), SARS-CoV-1 (P59594.1), bat_coronavirus_RaTG13 (QHR63300.2), pangolin_PCoV_GX-P5L (QIA48632.1), pangolin_PCoV_GX-P1E (QIA48623.1), bat_CoVZC45_CoV (AVP78031.1) and bat_Rs4874_CoV (ATO98205.1), were aligned using the T-coffee software (Rius et al., 2011). This multiple alignment was then used to estimate the phylogenetic relationships between SARS-CoV-2 and others coronaviruses. The phylogenetic tree was constructed using MrBayes (Huelsenbeck and Ronquist, 2001) with the following parameters: Likelihood model (Number of substitution types: 6(GTR); Substitution model: Poisson; Rates variation across sites: Invariable + gamma); Markov Chain Monte Carlo parameters (number of generations: 100 000; Sample a tree every: 1000 generations) and Discard first 500 trees sampled (burnin).

## ACKNOWLEDGMENTS

I would like to thank the IBIS bioinformatics group for their assistance. I am grateful to Dr. Michel Cusson, Université Laval, for the revision of the manuscript.

## CONFLICT OF INTERESTED

The author declares that he has no conflicts of interest.

